# The neural and computational architecture of feedback dynamics in mouse cortex during stimulus report

**DOI:** 10.1101/2023.07.19.549692

**Authors:** Simone Ciceri, Matthijs Noude Lohuis, Vivi Rottschäfer, Cyriel MA Pennartz, Daniele Avitabile, Simon van Gaal, Umberto Olcese

**Author notes:** These authors contributed equally (Daniele Avitabile, Simon van Gaal, Umberto Olcese).

## Abstract

Conscious reportability of visual input is associated with a bimodal neural response in primary visual cortex (V1): an early-latency response coupled to stimulus features and a late-latency response coupled to stimulus report or detection. This late wave of activity, central to major theories of consciousness, is thought to be driven by prefrontal cortex (PFC), responsible for “igniting” it. Here we analyzed two electrophysiological studies in mice performing different stimulus detection tasks, and characterize neural activity profiles in three key cortical regions: V1, posterior parietal cortex (PPC) and PFC. We then developed a minimal network model, constrained by known connectivity between these regions, reproducing the spatio-temporal propagation of visual-and report-related activity. Remarkably, while PFC was indeed necessary to generate report-related activity in V1, this occurred only through the mediation of PPC. PPC, and not PFC, had the final veto in enabling the report-related late wave of V1 activity.

## Introduction

A long-standing objective in the investigation of the neural mechanisms of consciousness is to characterize the signatures of perceived compared to non-perceived sensory stimuli at the level of neurons and their interactions (Aru et al., 2012; Koch et al., 2016). A milestone in the study of sensory-evoked responses in the visual system has been the observation of a bimodal neural response in the primary visual cortex: an early-latency response coupled to stimulus presentation and a late-latency response that is only observed when agents report the detection of a visual stimulus (Del Cul et al., 2007; Supèr et al., 2001; van Vugt et al., 2018). In spite of an ongoing debate on the functional role of this late activity component in conscious perception (Cohen et al., 2020; Hatamimajoumerd et al., 2022; Koch et al., 2016; Sergent et al., 2021), there is agreement that it remains strongly correlated with conscious report. Furthermore, this hallmark of visual detection is preserved across species, and has been the subject of circuit-level investigations in both non-human primates (Supèr et al., 2001; van Vugt et al., 2018), ferrets (Yin et al., 2020) and mice (Allen et al., 2017; Oude Lohuis et al., 2022b; Steinmetz et al., 2019; Zatka-Haas et al., 2021).

The cortical origin of the late, report-related activity observed in V1 has been pinpointed to frontal areas. Experiments performed across species observed that correlates of sensory detection behavior (which also carry categorical information about the behavioral relevance of the detected stimuli) first originate in prefrontal areas and only later appear in association and sensory areas (Del Cul et al., 2007; Steinmetz et al., 2019; van Vugt et al., 2018; Yin et al., 2020). A recent study even demonstrated that activity in a secondary motor area (a cortical subdivision that, in mice, is considered to be part of the prefrontal cortex(Le Merre et al., 2021)) is necessary for this late activity to emerge in mice (Allen et al., 2017). However, it is currently not understood how late, report-related activity reaches sensory regions and whether, besides originating in prefrontal regions, it is also shaped by other cortical regions, and if so how. For instance, it is debated whether prefrontal regions directly trigger late, report-related activity in primary sensory cortices, or whether this is (also) mechanistically driven by intermediate regions, such as association areas in the parietal and temporal lobes (Fahrenfort et al., 2008; Fisch et al., 2009; Quiroga et al., 2008; Sikkens et al., 2019; van Vugt et al., 2018). Addressing this question is important to better characterize how patterns of cortical activity that have been linked to conscious report are generated and propagate through cortical regions, and is consequential for arbitrating between major theories of consciousness (COGITATE Consortium et al., 2023; Melloni et al., 2021; Seth and Bayne, 2022).

Nevertheless, it is currently unfeasible to fully dissect the circuit-level architecture underlying the origin and propagation of neural activity. Several options (chiefly optogenetics) are available to modulate the activity of individual cortical areas (Oude Lohuis et al., 2022b, 2022a, 2021), but this approach is unsuitable to causally manipulate individual connections between regions. On the other hand, projection-specific optogenetic inactivation is only moderately effective on synaptic terminals or has relatively low temporal dynamics (Rost et al., 2022). The alternative approach of silencing the activity of feedback-projecting neurons, while achieving high efficacy and fast temporal specificity, inevitably modifies the activity of source cortical regions as well (Huh et al., 2018; Tervo et al., 2016). For these reasons, we decided to develop a minimal model of neural dynamics (Chaudhuri et al., 2015; Joglekar et al., 2018), which allowed us to test the contribution of individual feedback pathways to generating and propagating report-related activity across the cortical network. Compared to previous studies following a similar approach for studying report-related activity in the human brain (Alilović et al., 2023; Castro et al., 2020; Dehaene et al., 2003; Dehaene and Changeux, 2005), we leveraged the recently established availability of functional and structural data in mice (Harris et al., 2019; Knox et al., 2019; Oude Lohuis et al., 2022b; Steinmetz et al., 2019) to develop a computational model with anatomically faithful connectivity strengths between cortical regions and capable of reproducing patterns of spiking activity observed in mice performing perceptual tasks. We developed a model network composed of mouse primary visual cortex (V1), posterior parietal cortex (PPC) and prefrontal cortex (PFC). We found that, while PFC is necessary to generate report-related activity in V1, this effect can only be exerted through association areas such as PPC, which determine its characteristics and has the final “veto” for initiating report-related activity in V1. Thus, an interplay between frontal and parietal cortical regions is required to effectively integrate neural correlates of perception with ongoing sensory-evoked activity.

## Results

### Detection of visual stimuli is coupled to large-scale activity patterns in dorsal cortex

We first aimed to replicate and expand earlier reports that visual detection in mice correlates with a bimodal response pattern in V1 (Oude Lohuis et al., 2022b) and with the emergence of report-related activity across multiple cortical regions (Allen et al., 2017; Pho et al., 2018; Steinmetz et al., 2019; van Vugt et al., 2018; Yin et al., 2020; Zatka-Haas et al., 2021). To this aim, we first analyzed neuronal activity collected in head-fixed mice performing an audio-visual change detection task (Oude Lohuis et al., 2022b, 2022a) (Fig. 1A). Mice were trained to report the change in the orientation of the presented visual stimulus, by performing – for instance – a left lick, and a change in the pitch of the presented auditory stimulus by performing a right lick (with contingencies counterbalanced across mice, see Materials and Methods for details). In this report we only focus on the processing of the visual stimuli. Multi-area laminar probe recordings were performed in the primary visual cortex (V1), posterior parietal cortex (PPC) and anterior cingulate cortex (ACC) (Fig. 1B). We computed stimulus-evoked spiking responses across the three areas as a function of the saliency of the visual stimulus (threshold or max change) and based on whether a stimulus was detected (hit trial) or not (miss trial). Trials from the max change condition will be referred to as “high saliency” and trials from the threshold condition as “low saliency” from here on. Of relevance, previous studies indicated that neuronal responses in both PPC and V1 did not show major deviations based on whether licking responses to full-field visual stimuli had to be done towards a detector positioned towards the left or right side of a mouse’s snout (Oude Lohuis et al., 2022b, 2022a).

**Figure 1:**
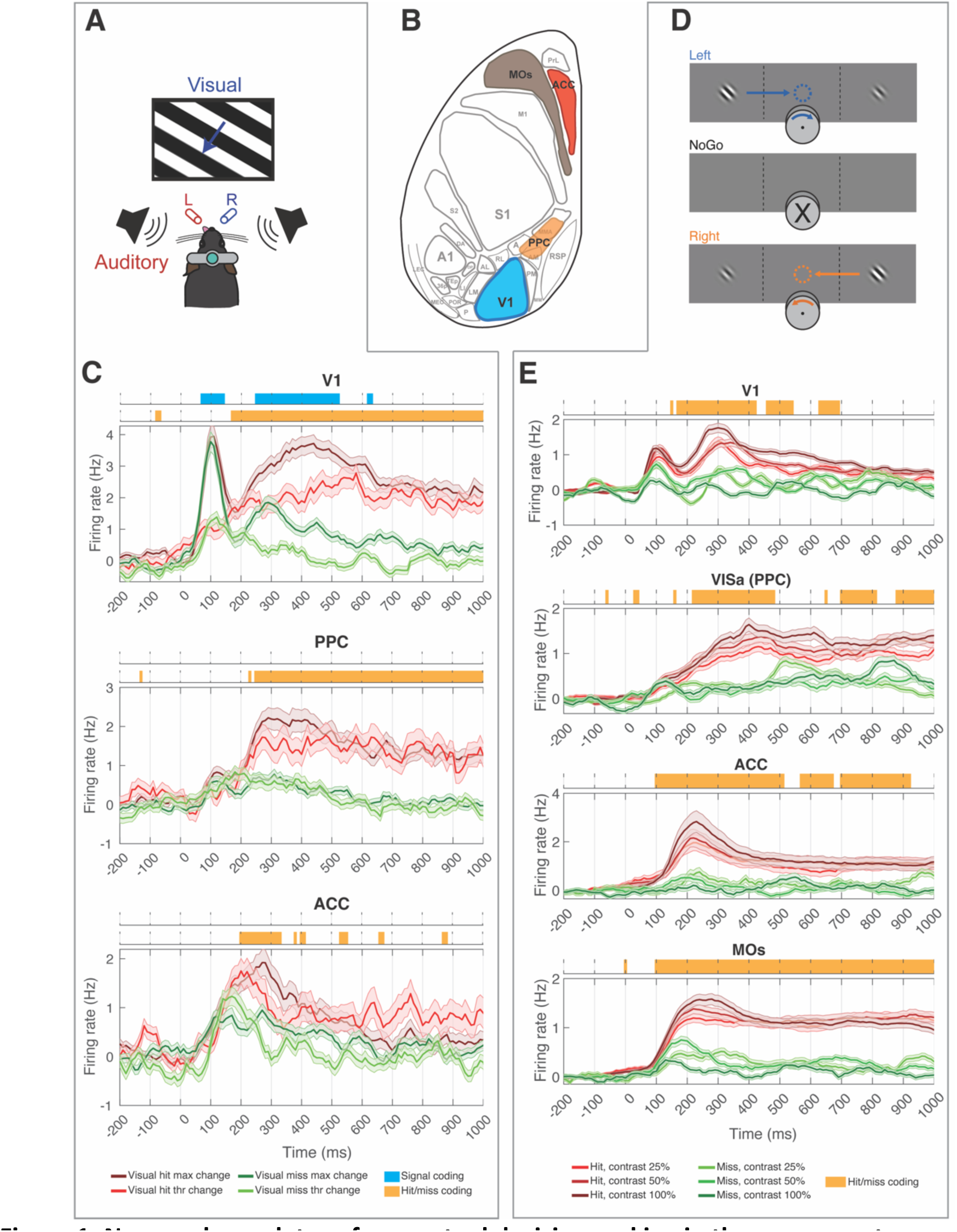
Neuronal correlates of perceptual decision making in the mouse cortex. (A) Schematic of the experimental configuration of the audio-visual change detection paradigm for head-fixed mice. Modified from ref. (Oude Lohuis et al., 2022a). (B) Schematic representation of the relevant cortical areas represented on a flattened cortical surface. Acronyms are used for the major subdivision of the dorsal cortex following standard nomenclature(Steinmetz et al., 2019; Wang et al., 2012; Wang and Burkhalter, 2007). Highlighted in color are the areas from which data was analyzed. V1: primary visual cortex; PPC: posterior parietal cortex; ACC: anterior cingulate cortex. MOs: Supplementary motor cortex. (C) Baseline-corrected average PSTHs recorded in (from top to bottom) V1, PPC and ACC following a change in the orientation of the presented drifting grating. Red: hits; green: misses. Dark colors indicate max visual change (highest saliency), while light colors indicate threshold visual change (low saliency). Shaded areas indicate the standard error of the mean. Color bars on top of individual panels indicate time bins in which significant differences (p<0.05, permutation-based test, FDR-corrected) were found between responses to – respectively – sensory stimuli with a difference salience (blue) or hit/miss trials (orange). (D) Outline of the contrast discrimination task, in which mice had to rotate a wheel to bring the Gabor patch with the highest contrast toward the center of the field of view. Modified from ref. (Steinmetz et al., 2019). (E) Same as C, but computed as a function of the difference in contrast between the stimulus presented in the contralateral field of view with respect to the recorded hemisphere (which was always the highest-contrast stimulus) and the stimulus presented ipsilaterally. The color darkness indicates the contrast difference. Statistical differences were computed as in panel C. Note that no difference between responses to sensory stimuli with different contrasts was observed.

In V1, we observed a bimodal pattern of activity: an early-onset wave of sensory-evoked activity, lasting until about 200 ms after stimulus onset, followed by a late-onset wave which was mainly encoding whether a trial was a hit or miss (cf. (Oude Lohuis et al., 2022b), Fig. 1C). Early sensory-evoked activity did not differ between hit and miss trials, but firing rates were positively correlated with the saliency of visual stimuli (Fig. 1C). Instead, late activity encoded both whether a trial was a hit or miss, as well as whether the sensory input was strong or weak (Fig. 1C, cf. (Oude Lohuis et al., 2022b)). Activity in PPC and ACC mainly encoded differences between hit and miss trials, although a generalized increase in firing rates could be observed as a consequence of the presentation of sensory stimuli (Fig. 1C, cf. (Oude Lohuis et al., 2022a)).

To verify that these results were not specific to our experimental protocol, we also analyzed recordings from a previously published experiment (Steinmetz et al., 2019). In this paradigm, mice had to identify which of two visual stimuli presented in the right and left hemifield had the highest contrast, and rotate a wheel to move the highest-contrast sensory stimulus toward the center of the screen (Fig. 1D). We computed sensory-evoked responses as a function of both stimulus contrast (difference between the contrast of the two presented Gabor patches) and hit/miss responses, for trials in which the highest-contrast stimulus was shown in the hemifield of view contralateral to the recorded hemisphere. We analyzed neuronal responses in areas corresponding to those we also recorded [V1; VISa and VISam (not shown), which are two secondary visual cortices spatially overlapping with PPC; ACC], as well as in a region broadly defined as supplementary motor cortex (MOs), where report-related activity has been shown to originate (Allen et al., 2017) – Fig. 1B. Results were in line with those that we observed in our dataset. V1 showed a bimodal pattern of activity, with an early sensory-evoked response (that however, in contrast with our dataset, did not encode stimulus saliency) followed by a report-related bump in activity (Fig. 1E). Activity in higher-order regions followed the early response displayed in V1 and was only report-related. These results suggest that the spatiotemporal progression of visual- and report-related activity is mostly independent from the details of the task being performed.

Earlier findings (Allen et al., 2017; van Vugt et al., 2018; Yin et al., 2020) indicated that report-related activity showed an earliest peak in prefrontal regions, followed by PPC and V1. Our results suggest a similar picture for what pertains higher-order regions, with prefrontal areas (ACC, MOs) (Le Merre et al., 2021) showing earlier indications of hit/miss differences compared to PPC. The relative timing of the appearance of report-related activity in V1 is, however, less clear (cf. Fig. 1C and 1E), but is overall very close to that observed in prefrontal regions (Allen et al., 2017; van Vugt et al., 2018; Yin et al., 2020). Thus, while prefrontal regions remain the most likely candidate for the origin of report-related activity – as supported by the causal experiments performed by Allen et al. (2017) (Allen et al., 2017) – the mechanistic pathway via which this form of activity reaches other cortical areas remains unclear.

### A minimal network model reproduces the spatio-temporal propagation of visual- and report-related activity

In order to understand the possible network-level mechanisms underlying the spatiotemporal propagation of visual- and report-related activity across the cortical areas from which we recorded neuronal activity, we developed a minimal mean-field computational model of the cortical network that: *(i)* uses available connectomic data (Bressloff, 2014; Chaudhuri et al., 2015; Ermentrout and Cowan, 1980; Ermentrout and Terman, 2010; Joglekar et al., 2018) and *(ii)* is calibrated using *in vivo* recordings. For this reason, the model only includes the three cortical areas from which we performed *in vivo* neuronal recordings: V1, PPC and PFC (see Fig. 2A). The activity in each area is modeled with a firing-rate neural-mass model comprising one excitatory and one inhibitory population. Firing rate models of this type are a well-tested tool to describe macroscopic neuronal dynamics, as they average single-neuron spike rates (Bressloff, 2014; Chaudhuri et al., 2015; Ermentrout and Cowan, 1980; Ermentrout and Terman, 2010; Joglekar et al., 2018). Within each mass, the synaptic dynamic has a tunable dispersion time, and oscillatory dynamics are possible because of the coupling between the excitatory and inhibitory population (Coombes and Wedgwood, 2023). We also adopted a classical nonlinear sigmoidal firing rate for each neuronal population (see Materials and Methods for a complete description), which is standard for neural mass models in the literature (Bressloff, 2014; Chaudhuri et al., 2015; Ermentrout and Cowan, 1980; Ermentrout and Terman, 2010; Joglekar et al., 2018).

**Figure 2:**
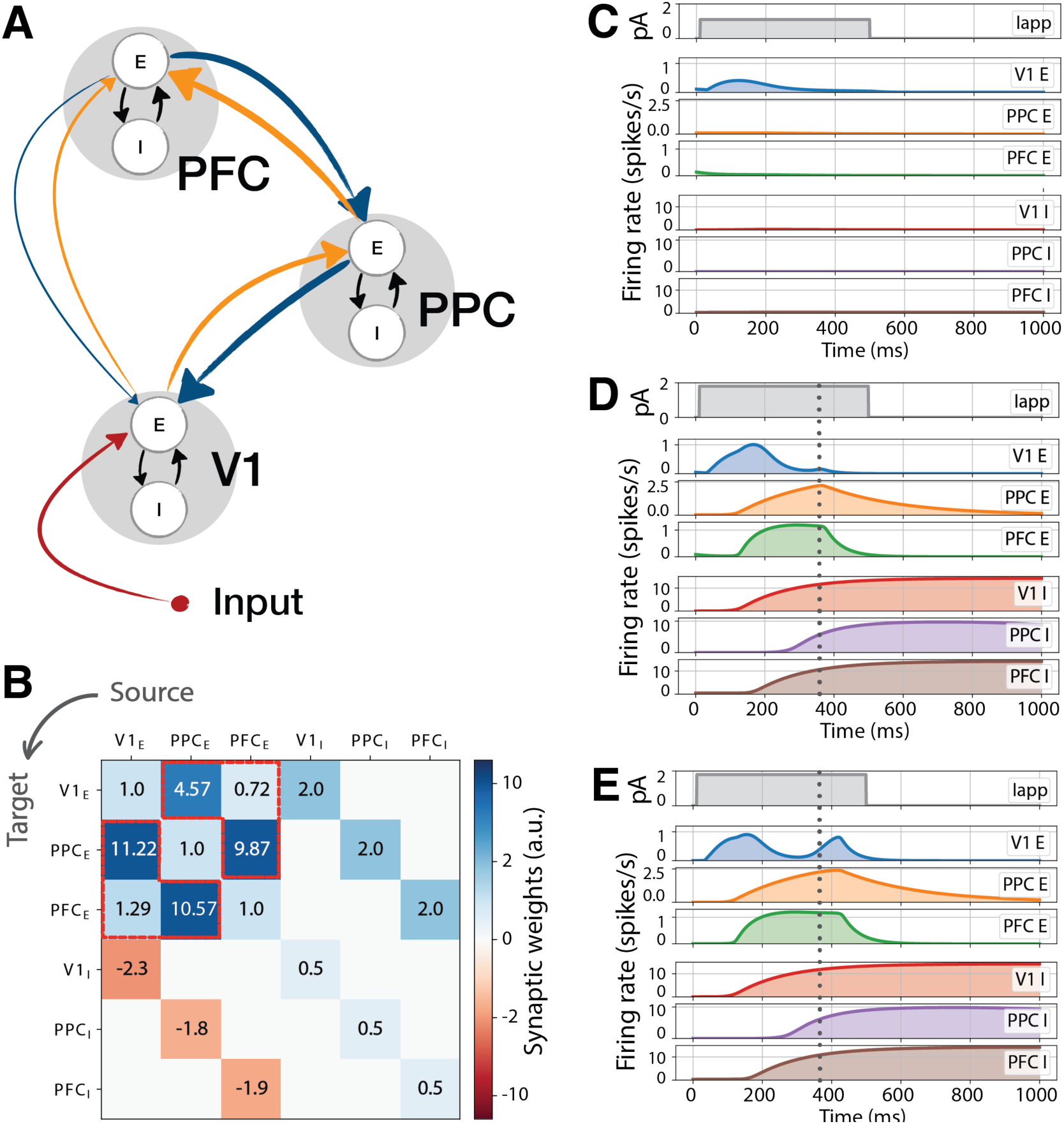
Network architecture and activity in the nominal setup. (A) Network schematic. We developed a minimal network model with an excitatory (E) and an inhibitory (I) node in three cortical areas: V1, PPC and PFC. Orange (blue) arrows indicate feedforward (feedback) connections, whose values were determined based on anatomical connectivity – roughly indicated by arrow thickness, see also panel *B*. Connections between excitatory and inhibitory nodes (black arrows) were calibrated to match experimental results. External input was applied to V1 (red arrow) to simulate visual stimuli. (B) Synaptic weights between nodes. The upper-left 3×3 block corresponds to non-local connections (matrix W, see STAR methods), while the other three blocks correspond to local couplings γ_i_. Values in highlighted cells (red lines) were experimentally derived. All other values were calibrated. (**C, D, E)** Example firing rate traces in the three regions for two different values of applied current: **C** low current I_max_ = 0.8pA; **D, E** medium current I_max_ = 2pA. At medium currents a feedback-bump may (*D*) or may not (*E*) appear depending on small changes in initial conditions. In *C-E*, row 1 reflects the input, rows 2-4 the activity of excitatory nodes and rows 5-7 the activity of inhibitory nodes.

The mean field model comprises a total of six neuronal populations, two in each cortical area, which feature local as well as long-range connections. More precisely, the excitatory-inhibitory pair in each cortical area are fully connected and should be taken together as a model of a cortical area, where the excitatory node has been fitted to experimentally collected neuronal activity and therefore represents the output of an area (in terms of “firing” activity) and all other variables represent hidden state variables. In addition, there are long-range excitatory connections to and from each cortical area. Crucially, connection strengths between areas were taken from recent experimental data (Knox et al., 2019) (see highlighted entries Fig. 2B). In particular, we employed values of directed connection density between V1, secondary visual areas A and AM (which are considered the mouse homologue of PPC (Arlt et al., 2022; Driscoll et al., 2017; Oude Lohuis et al., 2022a; Pinto et al., 2019)) and MOs (which is considered as a component of PFC (Le Merre et al., 2021) and is thought to be the key cortical area mainly in view of generating report-related activity (Allen et al., 2017; Steinmetz et al., 2019)). All other model parameters were calibrated (see Materials and Methods) to enable the excitatory nodes to reproduce patterns of activity comparable to those observed *in vivo*, as reported in earlier sections. During the tuning procedure all parameters in the model (characteristic rise/decay times, activation of the nonlinear firing rate functions, and local excitatory-inhibitory coupling strengths) were calibrated, while the inter-areal connections were kept fixed, because we had direct access to experimental data on these parameters. In this way we could test to what extent the generation and propagation of report-related activity in the three cortical areas on which we focus is shaped by cortico-cortical connectivity. Importantly, as our model comprises neural masses that jointly mimic the activity of whole cortical areas, all parameters except the activity of excitatory nodes do not represent measurable variables, but rather hidden state variables or input parameters that do not aim to model specific single-neuron parameters. For instance, while input currents are measured in picoampere, they reflect input to a whole cortical area and not to single neurons.

We modelled a visual stimulus via an applied transient step current, with varying intensity, on the excitatory population of V1 (see Fig. 2A, and *I_app_* time traces in Fig. 2C-E), and monitored the elicited cortical firing rate response in excitatory and inhibitory populations of V1, PPC, and PFC (whose time traces are also seen in Fig. 2C-E). To calibrate the model, we applied a visual input lasting for 500 ms to the excitatory V1 node and replicated the following experimental results. First, when subjected to a sufficiently strong stimulus, V1 activity displayed an early-onset response peaking around 100-200 ms (before the termination of the visual stimulus) that then dropped to lower values – Fig. 2C. This reproduces the adaptation to stimuli typically observed in the visual cortex (Fig. 1) – see e.g. (Kirchberger et al., 2021; Oude Lohuis et al., 2022b; Steinmetz et al., 2019; Supèr et al., 2001). Second, high-amplitude visual stimuli evoked a bimodal V1 response, that is, an early-onset peak of activity followed by a later peak – Fig. 2E. This second peak was absent if the stimulus had a low-amplitude (low salience stimulus) and accounts for the report-related activity observed *in vivo* (cf. hit trials in Fig. 1). While the model itself does not include an actuator stage to perform an actual report, we consider the emergence of the late-activity bump to represent an instance of stimulus detection by prefrontal/premotor cortical regions. This mimics the spatiotemporal time course of sensory detection, as can be observed from the neural recordings shown in Fig. 1 and from the related publications (Oude Lohuis et al., 2022b; Steinmetz et al., 2019).

Following this tuning procedure we observed that, when the network was subjected to stimuli of various intensity for 500ms, it displayed a strongly nonlinear response for the late peak of report-related activity: for smaller inputs (𝐼_max_ = 1.1𝑝𝐴) V1 was activated, but the signal did not significantly propagate in the network and did not trigger a second, feedback-dependent bump in activity (see excitatory nodes in row 2-4 of Fig. 2C). For higher input strengths (𝐼_max_ = 2𝑝𝐴), the network displayed the late activity bump in many, but not all, realizations – cf. Fig. 2D,E. To run the model, we first set the initial value of the firing rate in each of the excitatory and inhibitory population, that is, we set initial conditions prior the initiation of the stimulus. In the simulation each population is set to a random initial state with a small variance. In particular, we found that initial conditions varying within 10^-2^ spikes/s, simulating noisy initial data, trigger or suppress the occurrence of the late activity bump.

This is in line with experimental findings showing that, when subject to a sufficiently large stimulus, large late-latency activity arises with high probability, but not with certainty (van Vugt et al., 2018). The probabilistic nature of this response is analysed in detail in later sections. Before addressing this aspect, we observed the dynamics in each neuronal population, and noted that the occurrence of a late-latency activity bump appears to be feedback-induced. The external stimulus activated V1 which, in turn, following a feed-forward chain, activated PPC and PFC. The activity in the latter areas reached a peak (indicated with a vertical time-marker in Fig. 2D-E) before decaying owing to local (intra-population) inhibition. The time marker aligned remarkably well with the small late-latency activity in V1, signaling the onset of a feedback mechanism (from PFC and PPC back to V1, see schematic in Fig. 2A). In realizations in which the late activity bump occurred, it was again PFC and PPC that displayed a peak preceding the late, report-related activity in V1, in line with a feedback mechanism. This claim will be further substantiated in the following sections.

Thus the model, using a set of nominal parameters, was able to qualitatively reproduce the types of activity we observed *in vivo*. While the model was specifically tuned to reproduce V1 activity, we also obtained comparable patterns of activity in PPC and PFC, indicating that the model could be used to study the mechanisms underlying the propagation of activity across cortical areas during sensory-motor transformations. In particular, we focused on studying the role of feedback connections in the genesis of the late activity bump, i.e. of report-related activity.

We highlight that the time courses of the activity of excitatory nodes are strongly determined by the inhibitory ones: in Fig 2D-E it is visible that inhibitory nodes in each cortical area activate after the corresponding excitatory node, and this determines the rise-and-fall behavior in the latter. It is known that neurotransmitter release in excitatory and inhibitory populations are affected by time-scale separation between signals (Rodrigues et al., 2016). Our model achieves the delayed inhibitory activation using time-scale separation between excitatory and inhibitory rising times (see discrepancies in the parameters 𝜏^$^and 𝜏^%^in Materials and Methods). However, we also must point out that, for the purposes of this study, we did not calibrate the dynamics of inhibitory nodes to match experimental results, but only tuned their parameters so that excitatory nodes would show realistic behaviors. For this reason, we will only focus on excitatory nodes in the rest of the manuscript.

### The likelihood of late activity bumps is influenced by variations in internal state

As we have seen above, when the network is in the nominal setup and the visual stimulus is sufficiently high, a late activity bump occurs with a given probability, upon perturbing the initial state of the system. We investigated systematically this scenario by running 100 simulations during which the initial state of the excitatory V1 population was picked randomly and uniformly between 0 and 0.05 spikes/s, thereby imposing a small variance in the internal state (see Fig. 3A) that is in line with experimental observations linking cortical state fluctuations to perception (McGinley et al., 2015a, 2015b; Samaha et al., 2020; Speed et al., 2019; Supèr et al., 2003). We observed that early V1 responses elicited by either small (𝐼_max_ = 1𝑝𝐴) or large (𝐼_max_ = 3𝑝𝐴) input currents were not affected by such small variations, as trajectories were grouped together as simulation time progressed. On the other hand, intermediate currents (𝐼_max_ = 1.8 − 2𝑝𝐴) considerably propagated the initial uncertainty: a late activity bump occurred often, but the fine details of the trajectory could differ. These findings further support the conclusion that the network in the nominal setup supports robustly self-generated late-latency activity bumps. However, more delicate questions arise: given a fixed set of network parameters, how often does the network generate such a bump? And further: how do changes in the network parameters affect this likelihood? To address these questions, we developed first a mathematical index to track late-latency V1 activity.

**Figure 3:**
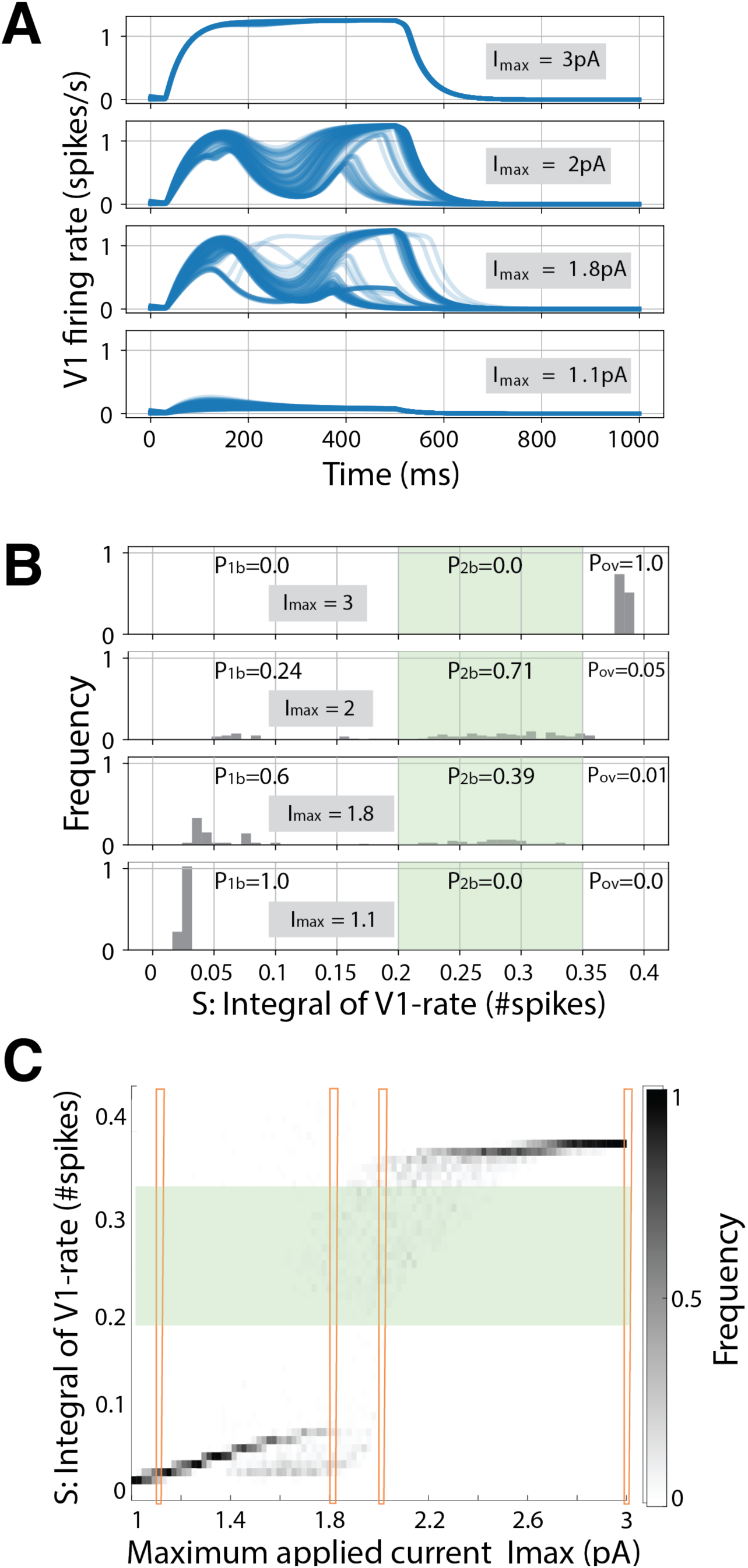
Network activity as a function of initial conditions and input currents. (A) Examples of activity trajectories (random initial conditions) for different input currents, in the nominal setup of parameters. (B) Distributions of integral quantity *S* for the trajectories in *A*. A late-activity bump is detected when the integral of V1 firing rate *S* lies in the interval [0.2, 0.35], highlighted in green. (C) Frequency distribution of *S* for different values of applied current I_app_. Red columns highlight the values displayed in *B*. The statistics is obtained over 100 different realizations for every value of I_max_, with initial conditions sampled from a uniform, random distribution u_i_(t = 0) ∈ [0, 0.05] spikes/s.

We therefore introduced a cumulative (integral) spiking measure *S*, with the view of determining the likelihood of late-latency activity in V1. For each of the V1 traces seen in Fig. 3A, we counted the average cumulative number of spikes occurring after the early activity bump in V1 (Materials and Methods). More precisely, we disregarded the trajectory before the reference time t_init_ = 250ms, because this is the characteristic time in which the early activity bump occurs (Del Cul et al., 2007; Oude Lohuis et al., 2022b), and then we calculated the area under the curve (proportional to the average number of population spikes) between 𝑡_*init*_ and the end of the simulation, 𝑡_,*-_= 1000ms, during which the late activity bump may occur. We expected trajectories with 𝐼_max_ = 1.1𝑝𝐴 (see Fig. 3A) to have a very small cumulative spike number *S*, because they did not display a late activity bump. Indeed, the histogram in Fig. 3B with 𝐼_max_ = 1.1𝑝𝐴 shows that in all such trajectories fewer than 0.05 spikes were observed in V1, on average, after the early activity bump, in the time interval [250ms, 1000ms]. On the contrary, a fully saturated response, in which firing rate reach the maximum value allowed by the model’s equations (Fig. 3A, 𝐼_max_ = 3𝑝𝐴), is characterized by a large *S*, and indeed the histogram in Fig 3B (with 𝐼_max_ = 3𝑝𝐴) shows that all such trajectories had more than 0.35 spikes after the first bump, on average. Finally, a late activity bump was characterized by an intermediate value of *S*: with 𝐼_max_ = 1.8 − 2𝑝𝐴 we observed a clear separation in the histogram of *S*. Therefore, we can use the value of *S* to define, empirically, the occurrence of a late activity bump (Fig. 3B). We thus classified a V1 activity trace by the corresponding value of *S*: we labeled traces with 0 < 𝑆 < 0.2, as displaying only the early activity bump, traces with 0.2 ≤ 𝑆 ≤ 0.35 (green band in Fig. 3B and 3C) as displaying both the early and late activity bumps and those with 𝑆 > 0.35 as displaying an overshoot (see also Fig. 3A).

Based on the fraction of trajectories whose S value falls in each of these three bands, we estimated the probability of having only the early activity bump as *P_1b_*, both the early and late activity bumps as *P_2b_* and overshoot as *P_ov_*. For example, from the histograms in Fig. 3B with stimulus 𝐼_.//_ = 2pA, we estimated that the nominal network displays both an early and late activity bump with probability 𝑃_01_ = 71%, an overshoot with probability 𝑃_23_ = 5%, and an early activity bump only or inactivity with probability 𝑃_41_ = 24%.

### Late-latency activity relies on network feedback

Armed with a quantitative index to inspect the likelihood of late-latency activity, we investigated how this likelihood changes upon variations in the network topology. As we shall see below, this analysis revealed that the late-latency activity is feedback-induced. We performed two different experiments which transform the connectivity matrix. In the first one, we perturbed the connectivity matrix at the initial time, and kept the matrix constant thereafter. In the second experiment we dynamically perturbed the matrix to explore the effects of abrupt changes to the topology of the network.

With the view of imposing changes in the network connectivity, we introduced a morphing parameter 𝛼 (see Materials and Methods). When 𝛼 = 1, the network is in its nominal state (the one studied so far); when 𝛼 > 1, selected network links are strengthened; when 𝛼 < 1, those links are weakened; finally, when 𝛼 = 0 the links are absent. The name morphing parameter suggests that with this index we can continuously transform the nominal network to intensify weaken, or even suppress certain links. Therefore, we introduced a tool to causally study to what extent the specific strength of an inter-area connection enables the emergence of a regime in which a sensory input to V1 can (with a certain probability) determine the occurrence of a late, report-related bump in activity.

We first used the morphing parameter to vary the strength of selected networks links from the starting time onwards, beginning with the feedback link from PFC to PPC (Fig. 4A), signposted with a red arrow in the network schematic (mathematically, the PFCèPPC connection was scaled by factor 𝛼). We repeated the experiment of Fig. 3A-B for various values of the morphing parameter (𝛼 between 0 and 1.5) and recorded the probability of a single early bump 𝑃1_1_, both early and late bumps 𝑃_01_, and overshoot 𝑃_23_. We used 𝑃_01_to derive the heatmap showed in Fig. 4A: the lighter colors correspond to a higher probability of a late activity bump, while darker colors denote lower probability thereof. We also used isolines to indicate where the probability of a single bump 𝑃_41_ crosses 99% (green isoline) and where the probability of overshoot 𝑃_23_crosses 99% (orange isoline).

**Figure 4:**
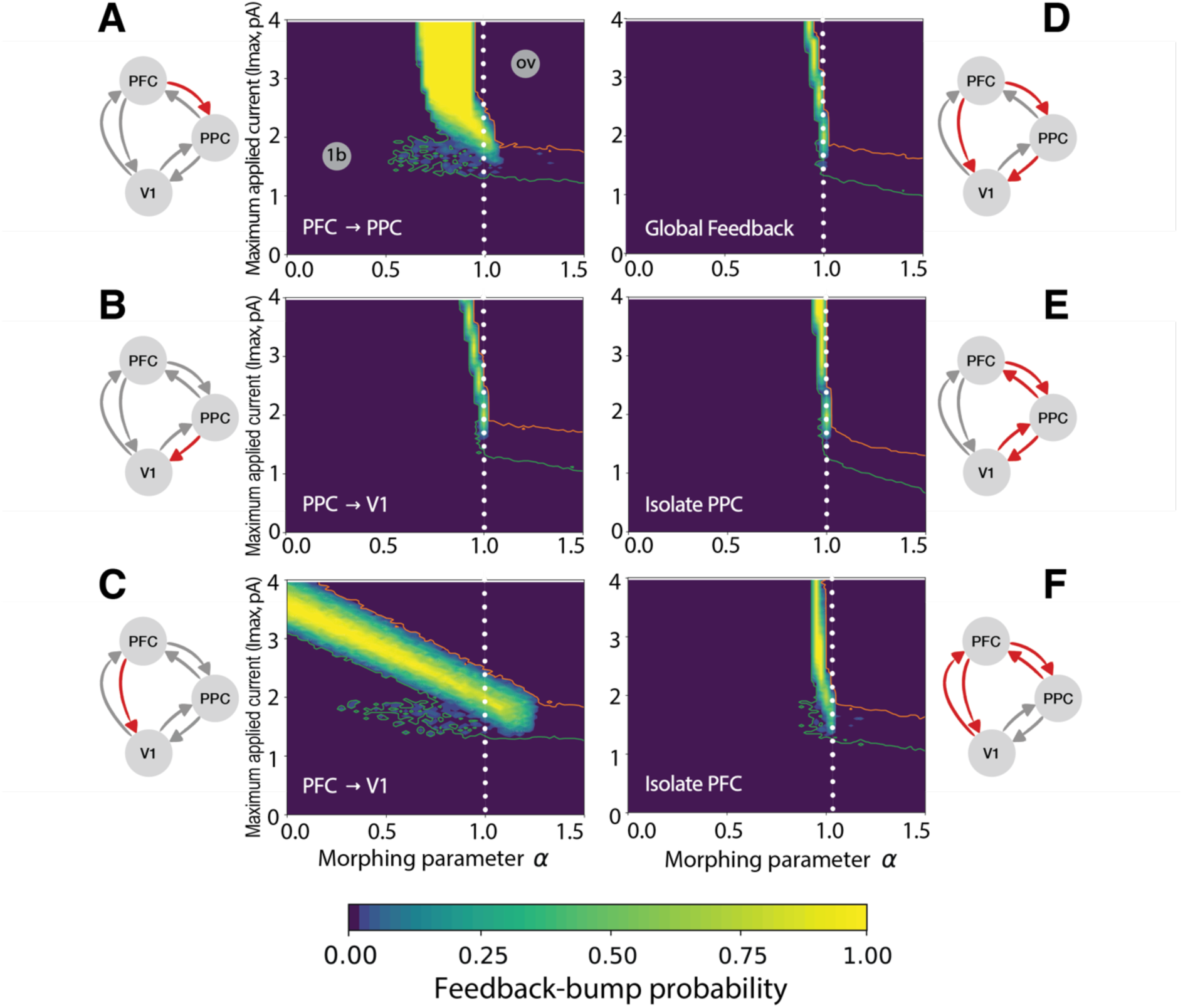
Probability of observing a late-activity bump, in the plane of parameters (α,I_max_), for the sets of connections considered. In each panel heatmaps show the probability of observing both an early and late activity bumps P_2b_ as a function of applied input current I_max_ and morphing parameter α, applied to a different set of connections. The green and orange lines represent isolines of P_1b_ = 99% and P_ov_ = 99% respectively, which border regions dominated by single bumps / inactivity and overshooting (see gray labels on top-left panel). A white, dotted line marks the nominal setup (α = 1). For each panel. morphed connections are colored in red on the respective network scheme. For each couple of (α,I_max_) values, we considered 50 random initial conditions (see Materials and Methods).

The overall information we gathered from the heatmap in Fig. 4A can be summarized as follows. First, the network produced a late activity bump robustly with respect to changes in the PFCèPPC feedback link: light yellow areas (late-bump probabilities close to 100%) were found in a variety of network configurations (for various values of 𝛼). Second, there are regimes, labelled “ov” and “1b”, where the feedback-bump was absent, but either an overshoot (ov) or an early bump only (1b) were found with probability greater than 99%, respectively. Third, the network could produce a late activity bump even when the feedback link PFCèPPC was weakened with respect to the nominal condition, provided that the strength of the impinging stimulus was increased; this can be deduced from the yellow area in Fig. 4A “curving upwards” towards higher values of 𝐼_max_. Finally, the feedback pathway PFCèPPC was an important player in triggering late-latency activity. While the network could compensate for the weakening of this link with a higher input to produce a late-latency bump, network configurations in which that link was either too weak or too strong failed to produce a late activity bump.

A markedly different behavior was observed when we perturbed the feedback link PPCèV1 (Fig. 4B). From Fig. 4B it can be seen that the likelihood of a late activity bump was strongly affected by changes in this feedback link. Small deviations from the nominal value of the link caused the late activity bump to disappear quickly. While the network could tolerate a weaker PFCèPPC link (Fig. 4A), even the slightest weakening of the PPCèV1 link caused a complete suppression of the feedback bump. This data revealed that the experimentally-derived anatomical connectivity used in the nominal conditions (𝛼 = 1) was crucial to obtain a late activity bump.

On the contrary, the network’s activity was only minimally affected by changes to the feedback link PFCèV1. Fig. 4C shows that feedback bumps could be produced with high probability even when this link was absent (𝛼 = 0), in case the strength of the input was increased (compensating for the reduced PFCèV1 link).

For the experiments in Figs. 4A-C we perturbed one link at a time, but the morphing parameter can also be varied on multiple links simultaneously. In Fig. 4D, for instance, we strengthened or weakened all the feedback pathways at once. These manipulations showed that late-latency bumps cannot exist without (or with too much) feedback. Therefore, it is the interplay between the various feedback pathways that generates the late-latency bump. This was further confirmed by the results displayed in Figs. 4E-F, showing that a network in which the PFC or PPC nodes were progressively isolated (transforming the architecture into a two-node network) did not display robust late activity bumps.

The previous experiment modulated the strength of feedback connections over the whole simulation period. However, we expect that, if the network receives a shock in the form of instantaneous removal of certain links during sensory processing (rather than from the initial time), this will also have an impact on late-latency activity. We investigated this scenario in a further experiment: we selected initial states which, in nominal conditions, would lead to late-latency activity; we then ran this network up to a chosen time T*, at which we instantaneously set selected links to zero (see Materials and Methods). In Fig. 5A.1 the PFCèV1 link is removed for different values of T* in the range 50-450 ms, when applying a value of I_max_=1.8pA. We found that inactivating the link at any time resulted in a quick decay in V1 activity, preventing the feedback bump to occur. In Fig. 5B.1 analogous results are found when PFC is instantaneously isolated from the rest of the network. We found similar results when we removed network links from the initial time (Fig. 4).

**Figure 5:**
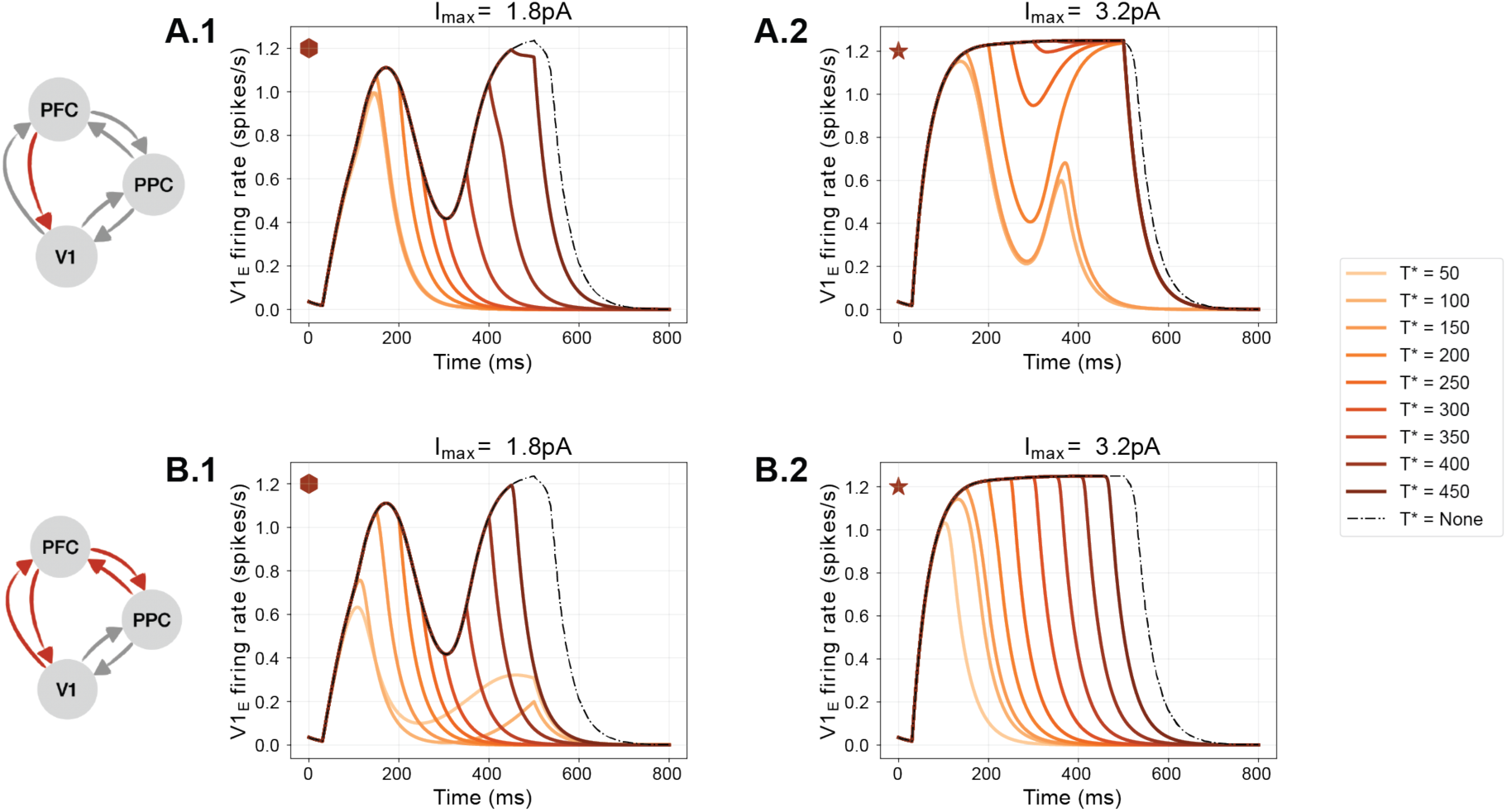
Effects of instantaneous link inactivation on feedback bumps. **(A)** We selected initial states and nominal conditions leading to late-latency activity (dashed line, T* = none). We ran this network up to a chosen time T*, and then instantaneously set the PFCèV1 link to zero. For I_max_=1.8pA (**A.1**) the instantaneous inactivation prevented (T*=50-300ms), or suppressed (T*>300ms) the feedback bump in V1. Conversely, when a higher value of I_max_ is used (A.2, I_max_=3.0pA), feedback activity can be restored in V1 despite the inactivation of the PFCèV1 link. (**B.1-2)** Same as A.1-2 but for the isolation of PFC from the network. Note how the feedback bump is always prevented or suppressed irrespective of the value of I_max_ (cf. Fig. 4, see also Fig. S1). The diamond and star shapes relate to the way the traces plotted in the panels can be retrieved in the heatmaps of Fig. S1.

However, as we discussed earlier, the amplitude of the external stimulus to V1 (i.e., I_max_) influences the genesis of the feedback bump. We thus repeated the previous experiments using a larger value of I_max_=3pA in order to establish whether sufficiently large inputs can compensate for instantaneous inactivation of the network links. In all cases, except for the PFCèV1 link, we found that large stimuli cannot overcome instantaneous inactivation of a set of links, hence impairing the generation of a feedback bump. In Fig. 5B.2, representative of most cases, it can be seen that a second bump is not formed, irrespective of the value T* (see also Fig. S1). On the contrary, a feedback bump is still visible when inactivation the PFCèV1 link with a T* between 150 and 300ms, showing that feedback activity can be restored by sufficiently large external drive in spite of the inactivation of this specific link.

In summary, the analyses performed using the morphing parameter shed light on the fact that the late activity bump is dependent on feedback connections. Upon external stimulation, V1 is activated and a feedforward pathway excites PPC and PFC. From these two areas, a late-latency activity bump appears in V1, mainly owing to an indirect pathway from PFC to PPC and then to V1. It turns out that direct feedback from PFC to V1 is instead not crucial for feedback bumps to be observed in V1, because the impact of reducing this link can be easily compensated by increasing stimulus strength.

## Discussion

In this project we set out to elucidate how report-related late activity patterns observed in cortex may mechanistically emerge as a function of the anatomically-constrained connectivity between sensory, association and prefrontal cortices.

We presented evidence of two crucial stages in the processing of visual information during perceptual decision making in mice, replicating and extending previous work in both humans and animals (Alilović et al., 2023; Allen et al., 2017; Dehaene and Changeux, 2011; Del Cul et al., 2007; Oude Lohuis et al., 2022b; Steinmetz et al., 2019; van Vugt et al., 2018). Neural recordings collected from two independent labs using two different tasks revealed that V1 firing rate at 100 ms after stimulus change was uniquely modulated by the saliency of the stimulus and not by the decision made by the mouse (Oude Lohuis et al., 2022b) (Fig. 1C,E). In contrast, a second, later wave of V1 activity reflected a combination of stimulus saliency and decision outcome. This late wave of activity was stronger for hits than for misses and coincided with increased report-related activity in both posterior parietal and frontal cortex.

Next, we set out to model the interactions between the three cortical regions that we considered, to understand the mechanisms giving rise to the observed neural dynamics. Crucially while designing the model connection strengths between V1, PPC and PFC were implemented from recent experimental data to match the currently known anatomical connectivity profiles between these regions. Thus, connectivity between cortical regions was imposed as a fixed constraint onto neural activity. Previous studies on the origin of report-related activity were either incorporating a limited set of cortical areas but did not constrain inter-area connectivity to experimentally-derived connectivity (Del Cul et al., 2007; van Vugt et al., 2018), or included most cortical regions but only globally varied inter-areal coupling strength (Castro et al., 2020; Dehaene et al., 2003; Dehaene and Changeux, 2005). Our approach allowed to mechanistically test the role of all combinations of single and multiple feedback connections with realistic strengths, something not feasible in whole-brain models. Importantly, thanks to the close link between experiments and our model, the time-specific inactivation experiment also provides predictions that, in contrast with more complex modeling approaches, are directly testable. Specifically, by optogenetically inactivating (i.e., isolating), PPC or PFC starting from different time points following stimulus onset (see e.g. (Kirchberger et al., 2021; Oude Lohuis et al., 2022b; Resulaj et al., 2018) for a similar approach) one could test the model validity in an existing experimental setup. Nevertheless, we cannot exclude that including additional areas known to play a role in top-down modulation of sensory areas (Halassa and Kastner, 2017; Wimmer et al., 2015; Zhang et al., 2016) might have partially modified the results we obtained. However, the fact that model behavior echoed the neural data observed *in vivo* indicates that the model’s architecture and the free model parameters were enabling physiologically-plausible sensory-evoked activity for a given, anatomically-based inter-areal connectivity matrix Nevertheless, it is important to briefly discuss the possible role of different areas. First, the anterior cingulate cortex (ACC) sends stronger feedback projections to V1 compared to other subdivisions of PFC such as MOs (Le Merre et al., 2021; Zhang et al., 2016). However, even though ACC has been shown to significantly modulate V1 activity (Fiser et al., 2016; Zhang et al., 2014), report-related activity has consistently been found to first appear in other PFC subdivisions compatible with MOs (Allen et al., 2017; Inagaki et al., 2022; Steinmetz et al., 2019; Takahashi et al., 2021; Yin et al., 2020). Second, while we focused on PPC, other temporal association areas have been implicated in visual perception (Conway, 2018). However, at least in rodents, medial association areas such as PPC, that have been hypothesized as the rodent homologue of the dorsal stream (Glickfeld et al., 2014; Wang et al., 2012), are more strongly connected to prefrontal areas compared to temporal association cortices (Harris et al., 2019; Knox et al., 2019). Furthermore, to what extent individual areas in the mouse association cortex (generally referred to as higher-order visual areas) provide unique contributions to sensory processing or form instead a redundant network is a matter of active debate (Glickfeld et al., 2014; Jin and Glickfeld, 2020; Keller et al., 2020; Kirchberger et al., 2021; Oude Lohuis et al., 2022a, 2021). Therefore, we believe that considering PPC as a single network node is an important first step to better understand the role of parietal and temporal cortex in the generation of report-related activity, to be followed-up by a more detailed characterization. Finally, although we cannot exclude a role of thalamic nuclei in mediating long-range top-down modulation, the thalamus seems to mainly play a modulatory role and not directly be involved in information transfer (Halassa and Kastner, 2017; Wimmer et al., 2015).

There were at least three notable observations in our model behavior. First, increasing the input strength to V1 led to both stronger early activity waves (∼100 ms) as well as an increased likelihood of a second late wave of activity (∼400 ms), as observed in our data and in previous studies (Del Cul et al., 2007; Supèr et al., 2001). Stronger V1 input also directly led to increased late activity in PPC and PFC, that in general anticipated the late wave of activity in timing observed in V1.

Second, we show that the initial baseline or pre-stimulus condition of each neural node strongly determines the likelihood of the late activity wave of activity to occur. In fact, we observed a regime of model parameters in which there was a clear nonlinear threshold for “igniting” this late wave of activity, in line with theoretical predictions from the Global workspace model of conscious access (Dehaene et al., 2003; Dehaene and Changeux, 2011, 2005; Mashour et al., 2020). This is in line with recent observations that variations in behavioral and cortical state, associated with ongoing fluctuations in pre-stimulus neural activity, strongly determine the likelihood that a stimulus will be reported (or consciously accessed) and that late report-related neural activity in cortex is observed (McCormick et al., 2020; McGinley et al., 2015a, 2015b; Speed et al., 2019; Waschke et al., 2019) .

Third, not only do we show that late activity in V1 is driven, as hypothesized earlier (Dehaene and Changeux, 2011), by feedback from higher-order regions, but also, more specifically, that although PFC is necessary to generate report-related activity in V1, it exerts its final influence on V1 only indirectly, through PPC. In our model, therefore, while PFC activity is necessary towards the buildup and initiation of the late, report-related bump in V1 activity, it is PPC that has the final veto and determines its characteristics. Removing the feedback connection from PFC to V1 had very limited influence on model behavior and the occurrence of a late activity wave could still be observed when stimulus input strength was increased. Thus, an interplay between frontal and parietal cortical regions is required, and most efficient, for eliciting late feedback activity in early sensory cortex (Supèr et al., 2001). This observation is important because, in the Global Neuronal Workspace Model, frontal cortex has always been considered the site responsible for igniting the network, and hence threshold setting (Dehaene, 2014) – but see (Sergent et al., 2021). It was hypothesized that once the threshold for conscious report in PFC is crossed, PFC sends information to other brain regions, including parietal cortex. We provide evidence for a possible different division of labor between frontal and parietal regions, in which frontal cortex acts as fast accumulator of sensory evidence, and parietal cortex as a much slower one (Kim and Shadlen, 1999; Shadlen and Newsome, 2001) – but see (Pinto et al., 2022). In this scenario, PFC starts to quickly feed stimulus evidence to parietal cortex, but activity in PFC quickly reaches a ceiling level. When activity in PFC reaches such a level, this is not sufficient to elicit a report-related wave of activity. In contrast, parietal cortex keeps on accumulating evidence from both sensory and prefrontal regions over time. Once activity in the PPC crosses a certain threshold (which can happen only if both V1 and PFC provide it with input activity), this triggers a report-related feedback wave of activity to V1 and other cortical areas. This scenario is a direct consequence of the known anatomically-constrained connectivity between areas (Harris et al., 2019; Knox et al., 2019), and specifically from the fact that PPC has a much stronger feedback connection to V1 compared to PFC.

In conclusion, our study provides a mechanistic hypothesis on the cortical pathway via which report-related activity, a hallmark of conscious access, is generated in the fronto-parietal cortex and then reaches sensory areas. Future experimental work will be required to validate our results *in vivo*. Furthermore, it will be important to disentangle potentially different mechanisms underlying the genesis of the different sub-components of late-onset activity in V1. In fact, recent studies suggest that this component of V1 activity correlates not only with report, but also with spontaneous and sensory-induced motor behavior (Lohuis et al., 2022; Oude Lohuis et al., 2022b; Steinmetz et al., 2019; Stringer et al., 2019). Finally, it is worth noting that the primary focus of our study lied in unraveling the mechanisms implicated in conscious access, associated with cognitive and behavioral responses to sensory stimuli (e.g., hits versus misses). Therefore, we did not aim to target the neural mechanisms of “phenomenal consciousness”: the subjective phenomenological aspects of conscious experience (Kriegel, 2007). In this light, it is rather uncontroversial that the fronto-parietal network is involved in conscious access or report, although exactly how so has been severely underspecified as we have highlighted before. The role of this network in phenomenal experience, and how this takes form in across the cortex, on the other hand, are strongly debated and a matter of ongoing investigation (Boly et al., 2017; Cohen et al., 2020; Hatamimajoumerd et al., 2022; Koch et al., 2016; Lamme, 2018; Odegaard et al., 2017). In fact, several theories of consciousness (e.g. integrated information theory (Tononi et al., 2016) or predictive processing accounts (Friston, 2010; Pennartz, 2022) – see (Seth and Bayne, 2022) for a review) emphasize the role of posterior association cortices (of which PPC is part) in the generation of phenomenal consciousness, and consider late activity to be only an reflection of conscious access, or of the subsequent motor actions. Thus, to what extent late, report-related activity reflects conscious processing, and what roles the different cortical regions implicated in its genesis play, remain open questions. A way to potentially investigate these issues is by employing so-called no-report paradigms (Tsuchiya et al., 2015), developed to isolate phenomenal experience from its functional consequences, such as report and decision-making, and pursue an interdisciplinary approach combining experiments and modeling. Whether this is possible and a fruitful approach, both theoretically as well as experimentally, is an important avenue for future scientific investigation.

## Materials and Methods

### RESOURCE AVAILABILITY

#### Lead contact

- Further information and requests for resources and reagents should be directed to and will be fulfilled by the lead contact, Umberto Olcese.

#### Materials availability

- This study did not generate new materials or reagents.

#### Data and code availability

- All the data collected by the authors of this study and used for the analyses presented in Fig. 1 will be shared by the lead contact upon request.
- This paper also analyzes existing, publicly available data. These accession numbers for the datasets are listed in the key resources table.
- All original code will be deposited at Zenodo and will publicly available as of the date of publication. DOIs will be listed in the key resources table.
- Any additional information required to reanalyze the data reported in this paper is available from the lead contact upon request.

### EXPERIMENTAL MODEL AND STUDY PARTICIPANT DETAILS

#### Collection and analysis of *in vivo* recordings

The model we developed (see *Model description*) was qualitatively fitted on data collected *in vivo* by the authors and on a previously released dataset (Steinmetz et al., 2019). The experimental procedures performed to collect the data are only summarized here, and are described more extensively in refs. (Oude Lohuis et al., 2022b, 2022a). All details about experimental subjects, recording procedures and behavioral task for the previously released dataset can be found in ref. (Steinmetz et al., 2019).

#### Experimental subjects

All animal experiments followed the relevant national and institutional regulations. Experimental procedures were approved by the Dutch Commission for Animal Experiments and by the Animal Welfare Body of the University of Amsterdam. The data presented here was collected from 17 male mice, obtained from two transgenic mouse lines: PVcre (B6;129P2-Pvalbtm1(cre)Arbr/J, RRID: IMSR_JAX:008069) and F1 offspring of this PVcre line and Ai9-TdTomato cre reporter mice (Gt(ROSA)26Sortm9(CAG-tdTomato)Hze RRID: ISMR_JAX 007909). Mice were group-housed in under a reversed day-night schedule (lights on at 20:00 and off at 8:00) and all experimental procedures were done in the dark period. Temperature was kept between 19.5 and 23.5 °C, and humidity between 45% and 65%. During behavioral training (starting when mice were about 8 week old), mice were kept under a water restriction regime. Their minimum weight was kept above 85% of their average weight between P60-P90. Mice were normally trained 5 days/week, and generally obtained all their daily liquids in the form of rewards during task performance. A supplement was delivered when the amount of liquid obtained during the task was below a minimum of 0.025 ml/g body weight per day. The same amount was provided during weekends. Mice received *ad libitum* food.

### METHODS

#### Surgical procedures

At the start of experimental procedures, mice were implanted with a headbar to allow head-fixation in the experimental setup. About three weeks before electrophysiological recordings, a subset of mice received an injection of an adeno-associated virus mediating the Cre-dependent expression of ChR2 in Parvalbumin-positive interneurons; the injection was performed, in separate sets of mice, in either V1 or PPC. Data collected during optogenetic interventions was not utilized for the analyses presented in this study. The day before the start of extracellular recordings, small craniotomies (about 200 μm in diameter) over the cortical areas of interest using a dental drill. Cortical regions (V1, PPC and anterior cingulate cortex ACC for this study) were identified either via stereotactic coordinates or via intrinsic optical signal imaging – Fig. 1A. Details about all surgical procedures can be found refs. (Oude Lohuis et al., 2022b, 2022a).

#### Behavioral task and sensory stimuli

Mice were trained, over the course of several weeks, to perform an audio-visual change detection task – Fig. 1C. Visual stimuli were drifting square-wave gratings (temporal frequency: 1.5 Hz; spatial frequency: 0.08 cycles per degree; contrast: 70%; gamma-corrected), presented over the full screen (18.5-inch monitor, 60 Hz refresh rate). Gratings were continuously presented at a distance of about 21 cm from the eyes. In a subset of trials (visual change trials) the orientation of the drifting grating was instantaneously changed. The amount of orientation change determined the visual saliency, which was set, based on the properties of the psychometric curve of individual mice, to a value corresponding to a threshold or max change (detection threshold and 90 deg, respectively). Mice were trained to respond to a visual change by licking to one reward port (left or right, counterbalanced across mice), and received 5-8 μl of liquid reward (infant formula) upon a correct response. Visual stimuli were the subject of analysis in the current manuscript, and a detailed description can be found in refs. (Oude Lohuis et al., 2022b, 2022a). Correct responses to an auditory changes corresponded to licks toward the port not rewarded for visual stimuli (counterbalanced across mice). Importantly, similar neuronal responses were obtained across the measured areas irrespective of the side to which the mice had to lick upon a visual change, as well as independently of whether mice were trained to only report visual but not auditory changes. A more in depth account can be found in refs. (Oude Lohuis et al., 2022b, 2022a).

#### Multi-area recordings: acquisition and pre-processing

Extracellular recordings were performed simultaneously in 2 or 3 cortical areas (V1, PPC, ACC and A1 were targeted in different experimental sessions). Recordings were performed on a maximum of 4 consecutive days. Several types of Neuronexus (Ann Arbor, MI) silicon probes were used (A1 × 32-Poly2–10 mm-50 s-177, A2 × 16-10 mm-100-500-177, A4 × 8-5 mm-100-200-177, A1 × 64-Poly2-6 mm-23 s-160). Neurophysiological signals were pre-amplified, bandpass filtered (0.1 Hz to 9 kHz), and acquired at 32 kHz (a band-pass filter was set between 0.1 Hz and 9 kHz) with a Digital Lynx SX 128 channel system, via the acquisition software Cheetah 5.0 (Neuralynx, Bozeman, MT). Spike detection and sorting were performed using the Klusta (version 3.0.16) and Phy (version 1.0.9) software packages. For more details about acquisition and pre-processing, refer to refs. (Oude Lohuis et al., 2022b, 2022a).

#### Histology

At the end of experiments, mice were perfused in 4% PFA in PBS, and their brains were recovered for histological reconstruction meant to verify the correct placement of silicon probes in V1, PPC and ACC.

#### Model description

We modeled a network of 3 regions, namely V1 (primary visual cortex), PPC (posterior parietal cortex) and PFC (prefrontal cortex). Each region comprises one excitatory and one inhibitory population (see schematic in Fig. 2), and the activity of each population is described by a neural mass model (Bressloff, 2014; Chaudhuri et al., 2015; Ermentrout and Cowan, 1980; Ermentrout and Terman, 2010; Joglekar et al., 2018). The model describes the evolution of the average population firing rates. Such models are macroscopic in nature, that is, they describe population activity, as opposed to single-neuron activity. Populations are connected through weighted links, which represent anatomical connectivities. Neural Mass Wilson-Cowan models, such as the ones described below, are an established framework to investigate large-scale neuronal dynamics (Bressloff, 2014; Chaudhuri et al., 2015; Ermentrout and Cowan, 1980; Ermentrout and Terman, 2010; Joglekar et al., 2018).

The 𝑖th cortical area in the network evolves according to the following equations:

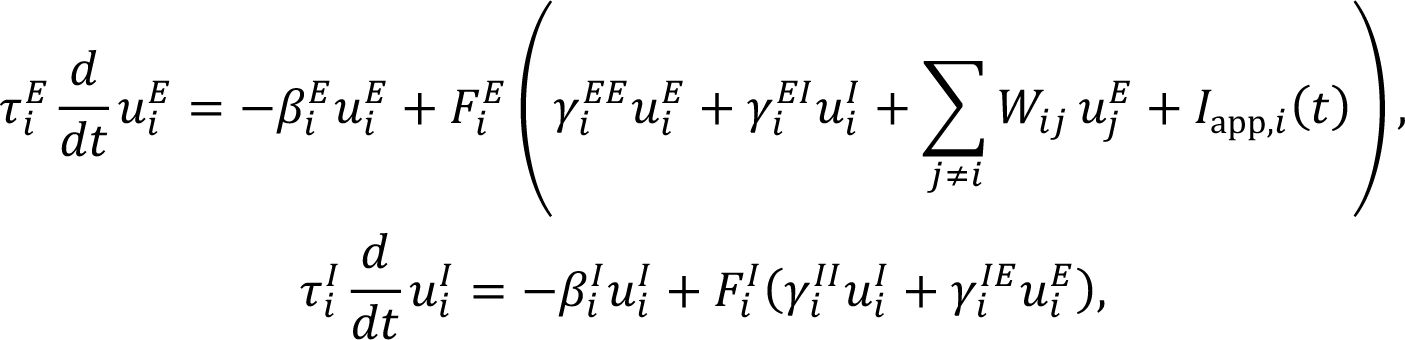

where the superscripts 𝐸, 𝐼 label excitatory and inhibitory variables, respectively. The firing rate 𝑢_*i*_ of the 𝑖-th population has characteristic time constants 𝜏_*i*_, and it evolves according to two main contributions: a damping term proportional to 𝛽_*i*_, and a nonlinear synaptic term, collecting inputs from the network. Our network is formed by 3 main brain regions (V1, PPC, and PFC) hence we set 𝑢_1_ = 𝑉1, 𝑢_2_ = 𝑃𝑃𝐶 and 𝑢_3_ = 𝑃𝐹𝐶, each endowed with an excitatory and inhibitory node, thereby obtaining a network with 6 nodes.

The local couplings are denoted by 𝛾*^kl^_i_* where 𝑘, 𝑙 = 𝐸, 𝐼. Inhibitory populations are connected only locally, whereas excitatory populations have local as well as long-range connections. Long-range connections are mutual, all-to-all and, in general, asymmetric. This means that, while each population is connected to all the others, the respective weights have different strengths. We encode the link from the excitatory population 𝑗 to the excitatory 𝑖 in a matrix using the equation:

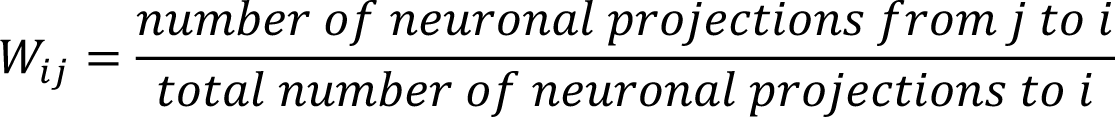

Nominal values of 𝑊_*ij*_ (see highlighted entries in the upper left 3×3 block of Fig. 2B) have been taken according to recent data on mice (Harris et al., 2019; Knox et al., 2019). This allows to develop a model with faithful connectivity between cortical regions.

The nonlinear function 𝐹^*E*^_*i*_ is sigmoidal:

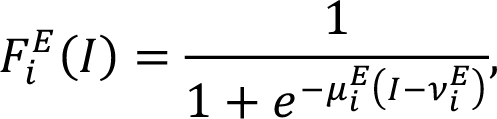

and a similar expression holds for 𝐹*^I^_i_*. The parameter 𝜇*^E^_i_* influences the sharpness of the sigmoid, while 𝜈*^E^_i_* determines the threshold at which the nonlinear firing response is triggered. Finally, we model a network receiving an external stimulus in V1, hence 𝐼_app_(𝑡) is different from 0 only in V1, so 𝐼_app,*i*_ = 0 for 𝑖 = 2,3 and it is a step function for 𝑉_1_

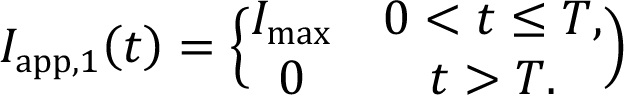

Nominal values of parameters are reported in the next paragraph. Parameter variations are discussed in the main text.

The described equations are numerically integrated using the function ode23s in Matlab, which is based on a modified Rosenbrock formula of order 2 (Shampine and Reichelt, 1997).

#### Numerical parameters values

Connection strengths between areas (matrix W) were taken from recent experimental data (Knox et al., 2019). All other model parameters were manually calibrated to enable the excitatory nodes to reproduce patterns of activity comparable to those observed *in vivo*, and reported earlier. As the model describes population activity, its variable 𝑢_*i*_ refers to the average firing rate of single neurons within population 𝑖. As customary in mean-field models, some parameters refer to single neurons, while others to entire population. For instance, the input current 𝐼_*app*_ is interpreted as the average external current received by a single neuron (measured in *pA*), and similarly for coupling and synaptic constants. On the other hand, characteristic timescales refer to populations.

|  | V1 | PPC | PFC |
| --- | --- | --- | --- |
| V1 |  | 11.22 | 1.29 |
| PPC | 4.57 |  | 10.57 |
| PFC | 0.72 | 9.78 |  |

*Gamma parameters (units: pA/Hz)*
|  | V1 | PCC | PFC |
| --- | --- | --- | --- |
| $\gamma^{EE}$ | 1. | 1. | 1. |
| $\gamma^{EI}$ | 2.3 | 1.8 | 1.9 |
| $\gamma^{IE}$ | 2 | 2 | 2 |
| $\gamma^{II}$ | 0.5 | 0.5 | 0.5 |

*Parameters of the firing rate function:*
|  | V1 | PPC | PFC |
| --- | --- | --- | --- |
| $\mu^E$ [1/pA] | 3 | 2 | 2 |
| $\mu^I$ [1/pA] | 2 | 2 | 2 |
| $\nu^E$ [pA] | 2 | 4 | 2 |
| $\nu^I$ [pA] | 0.3 | 0.3 | 0.3 |

*Characteristic times and decay constants:*
|  | V1 | PCC | PFC |
| --- | --- | --- | --- |
| $\tau^E$ [ms] | 30 | 200 | 38 |
| $\tau^I$ [ms] | 10 | 10 | 10 |
| $\beta^E$ | 0.8 | 0.9 | 3.8 |
| $\beta^I$ | 0.07 | 0.1 | 0.07 |

#### Noise in the initial conditions

We investigated the network behavior when the nominal setup is perturbed by noise, by sampling the initial conditions from a random, uniform distribution: u_i_(t=0) ∈ [0, 𝛿].

The analysis of network behavior under noise is obtained by with 50 realizations for each network setup. We fixed δ=0.05 spikes/s, as this value was one order of magnitude smaller than the typical scales of the excitatory firing rates, and we additionally studied the effects of varying 𝛿.

### Instantaneous Inactivation and connectivity morphing parameter

We perform two experiments to examine the robustness of the network behavior with respect to changes in the coupling between areas, and also to infer which nodes are most relevant for the formation of the late activity bump. First, we introduce a connectivity morphing parameter which amplifies (α>1), dampens (or suppresses (α<1) one or more synaptic connections (*W_ij_*) with respect to their nominal value (α=1), via the following transformation:

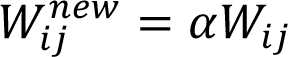

This transformation is performed at the initial time, and allows for intermediate states of weakened connections In a second experiment we consider instantaneous inactivation of certain synaptic connection: at specific time T*, we set one or more entries of the connectivity matrix (𝑊_*ij*_) to zero:

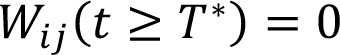

### QUANTIFICATION AND STATISTICAL ANALYSIS

#### In vivo recordings: Data analysis

All data analysis was performed in Matlab 2021b (MathWorks).

#### In vivo recordings: Sensory-evoked and task-related responses

For each single neuron identified through the spike sorting procedure (V1: 594 neurons, PPC: 529 neurons, ACC: 629 neurons), we computed the average peri-stimulus time histogram (PSTH) aligned to the onset of visual changes, separately for hit and miss trials, as well as for small and large visual changes. PSTHs were computed with a 10 ms time bin, and smoothed with a Gaussian window (standard deviation: 25 ms). Each PSTH was baseline-corrected, i.e., we subtracted the average activity computed in the [-500 -10] ms window with respect to stimulus onset.

#### In vivo recordings: Sensory-evoked and task-related responses – previously released dataset

The dataset used for the Steinmetz et al. (2019) (Steinmetz et al., 2019) study was downloaded from https://figshare.com/articles/dataset/Dataset_from_Steinmetz_et_al_2019/9598406 and analyzed using the same approach described above. Trials were pooled together based on whether a hit or miss was observed, and separately for visual contrasts of 25%, 50% and 100%.

#### In vivo recordings: Statistical analyses

Differences between sensory-evoked responses were assessed using a permutation-based approach. For each pair of conditions to be tested (e.g. hit trials to high vs. low saliency stimuli) we used the corresponding single-neuron PSTHs to compute the difference between average responses (across neurons) separately for each time bin. We then randomly swapped the trial identify of each PSTH, separately for each neuron, and computed the corresponding response difference. This was repeated 1000 times. We then ranked, separately per time bin, the actual response difference between two conditions compared to the values obtained through random permutations. If the actual response difference was higher than 95% of the values obtained through random permutations, a difference was considered to be significant, and the corresponding p value was compute as the fraction of randomly obtained values which was higher than the actual difference. All p values were then corrected for the false discovery rate (Bonferroni correction). To compute if an area encoded differences between high and low saliency stimuli, we further specified that this difference had to be present for both hit and miss trials, to prevent any interaction effect. Similarly, any hit/miss difference had to be present for both low and high saliency stimuli.

#### Model: Neural activity measure

We define an integral measure which counts the cumulative number of spikes in V1, from time *t_init_* to time *t_end_*

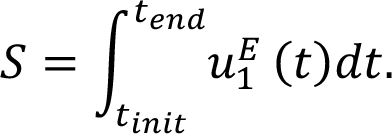

Here 𝑢^*E*^_1_(𝑡) is the firing rate of the excitatory population in the primary visual cortex 𝑉_1_. We shall set 𝑡_*init*_ so as to start counting spikes after a first (stimulus-induced) bump occurs, and use 𝑆 to determine whether a second (feedback-induced) bump is present in V1.

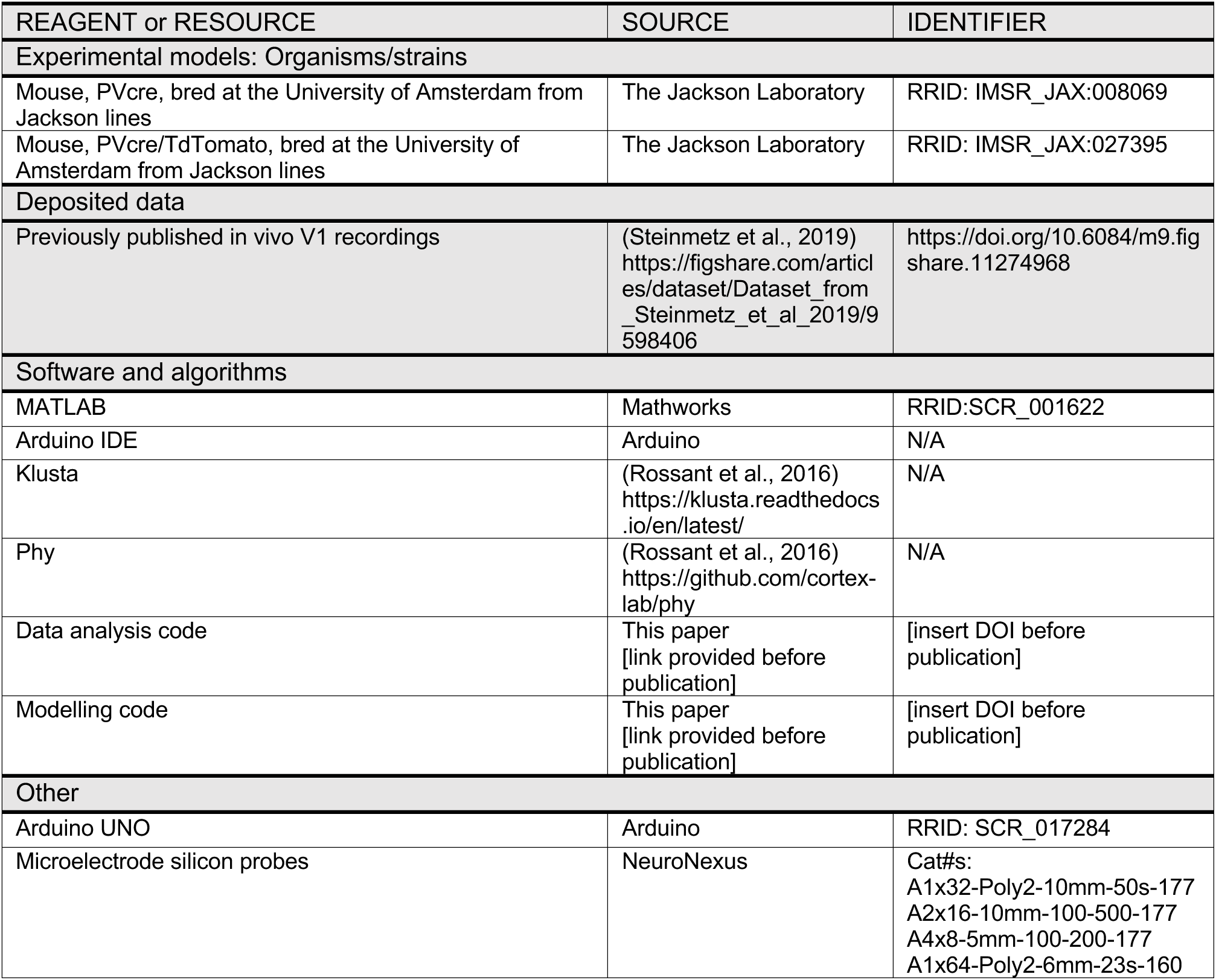
KEY RESOURCES TABLE

KEY RESOURCES TABLE
| REAGENT or RESOURCE | SOURCE | IDENTIFIER |
| --- | --- | --- |
| Experimental models: Organisms/strains |  |  |
| Mouse, PVcre, bred at the University of Amsterdam from Jackson lines | The Jackson Laboratory | RRID: IMSR_JAX:008069 |
| Mouse, PVcre/TdTomato, bred at the University of Amsterdam from Jackson lines | The Jackson Laboratory | RRID: IMSR_JAX:027395 |
| Deposited data |  |  |
| Previously published in vivo V1 recordings | (Steinmetz et al., 2019)<br>https://figshare.com/articles/dataset/Dataset_from_Steinmetz_et_al_2019/9598406 | https://doi.org/10.6084/m9.figshare.11274968 |
| Software and algorithms |  |  |
| MATLAB | Mathworks | RRID:SCR_001622 |
| Arduino IDE | Arduino | N/A |
| Klusta | (Rossant et al., 2016)<br>https://klusta.readthedocs.io/en/latest/ | N/A |
| Phy | (Rossant et al., 2016)<br>https://github.com/cortex-lab/phy | N/A |
| Data analysis code | This paper<br>[link provided before publication] | [insert DOI before publication] |
| Modelling code | This paper<br>[link provided before publication] | [insert DOI before publication] |
| Other |  |  |
| Arduino UNO | Arduino | RRID: SCR_017284 |
| Microelectrode silicon probes | NeuroNexus | Cat#s:<br>A1x32-Poly2-10mm-50s-177<br>A2x16-10mm-100-500-177<br>A4x8-5mm-100-200-177<br>A1x64-Poly2-6mm-23s-160 |

## Acknowledgments

This work was supported by Netherlands Organization for Scientific Research (NWO Crossover project INTENSE to U.O. and C.M.A.P.), the European Research Council (ERC Starting grant, 715605 project CONSCIOUSNESS to SVG). This project was also made possible through the support of a grant from Templeton World Charity Foundation, Inc. (funder DOI 501100011730) through grants TWCF0646 (to U.O. and C.M.A.P.) and TWCF-2022-30261 (to U.O.). The opinions expressed in this publication are those of the authors and do not necessarily reflect the views of Templeton World Charity Foundation, Inc. The authors would also like to thank Eric Dijkema for providing support in accessing the data used for Fig. 1, and the Carandini/Harris lab for providing free access to the raw data used in ref. (Steinmetz et al., 2019). The authors would also like to thank the Institute for Advanced Studies of the University of Amsterdam for supporting a workshop that spearheaded this study.

## Author contributions

Conceptualization: VR, DA, SvG, UO; Methodology: VR, DA, SvG, UO; Software: SC, DA, UO; Formal Analysis: SC, DA, UO; Investigation: SC, DA, MoL; Resources: MNoL; Data Curation: MoL; Writing – Original Draft: SC, VR, DA, SvG, UO; Writing – Review & Editing: SC, MNoL, CMAP, VR, DA, SvG, UO; Visualization: SC, DA, UO; Supervision: DA, SvG, UO; Funding Acquisition: CMAP, DA, SvG, UO.(Steinmetz et al., 2019)

## Supplementary figures

**Figure S1:**
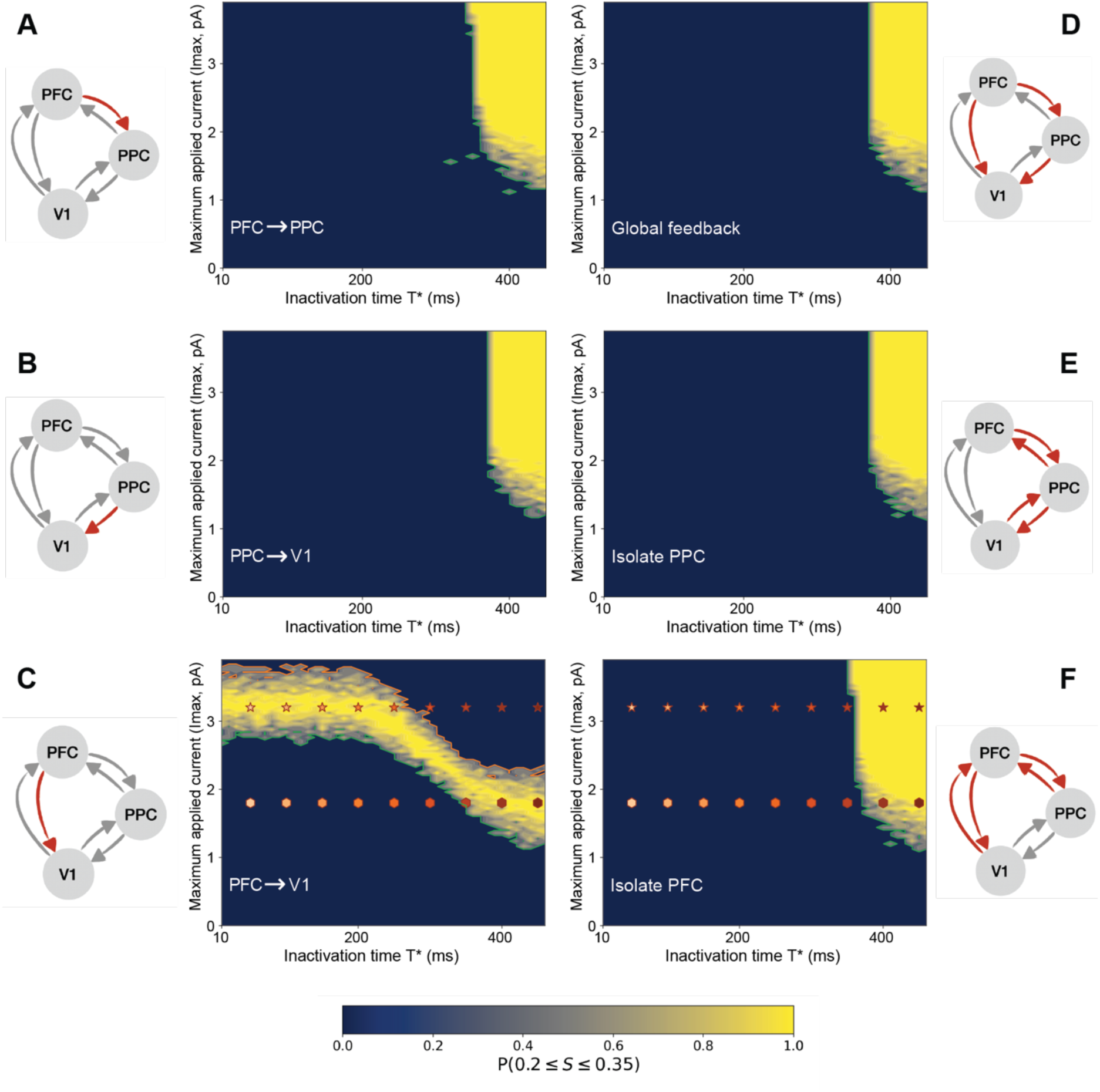
Probability of observing a late-activity bump, in the plane of parameters (T*,I_max_), for the sets of connections considered. In each panel (same order as Fig. 4) the heatmaps show the probability of the measure S indicating a feedback bump (0.2 ≤ 𝑆 ≤ 0.35, as a function of stimulus strength I_max_ and the instantaneous inactivation time T*, applied to a different set of connections. The green and orange lines represent isolines of probabilities P(S<0.2) = 99% and P(S>0.35) = 99%, respectively. For each panel, the morphed set of connections is colored in red in the respective network scheme. For each pair of (T*,I_max_) values, we considered 50 random initial conditions. Early link inactivation (T*<300) prevents the formation of the feedback bump for all chosen sets of connections, with the exception of PFCèV1 (panel C). The diamond and star shapes in panels C and F indicate the pair of (T*,I_max_) values for which we plotted V1 dynamics in Fig. 5 (diamonds: I_max_=1.8pA; stars: I_max_= 3.0pA).

