## Supplementary figures and images for "The neural and computational architecture of feedback dynamics in mouse cortex during stimulus report"

### Figure S1

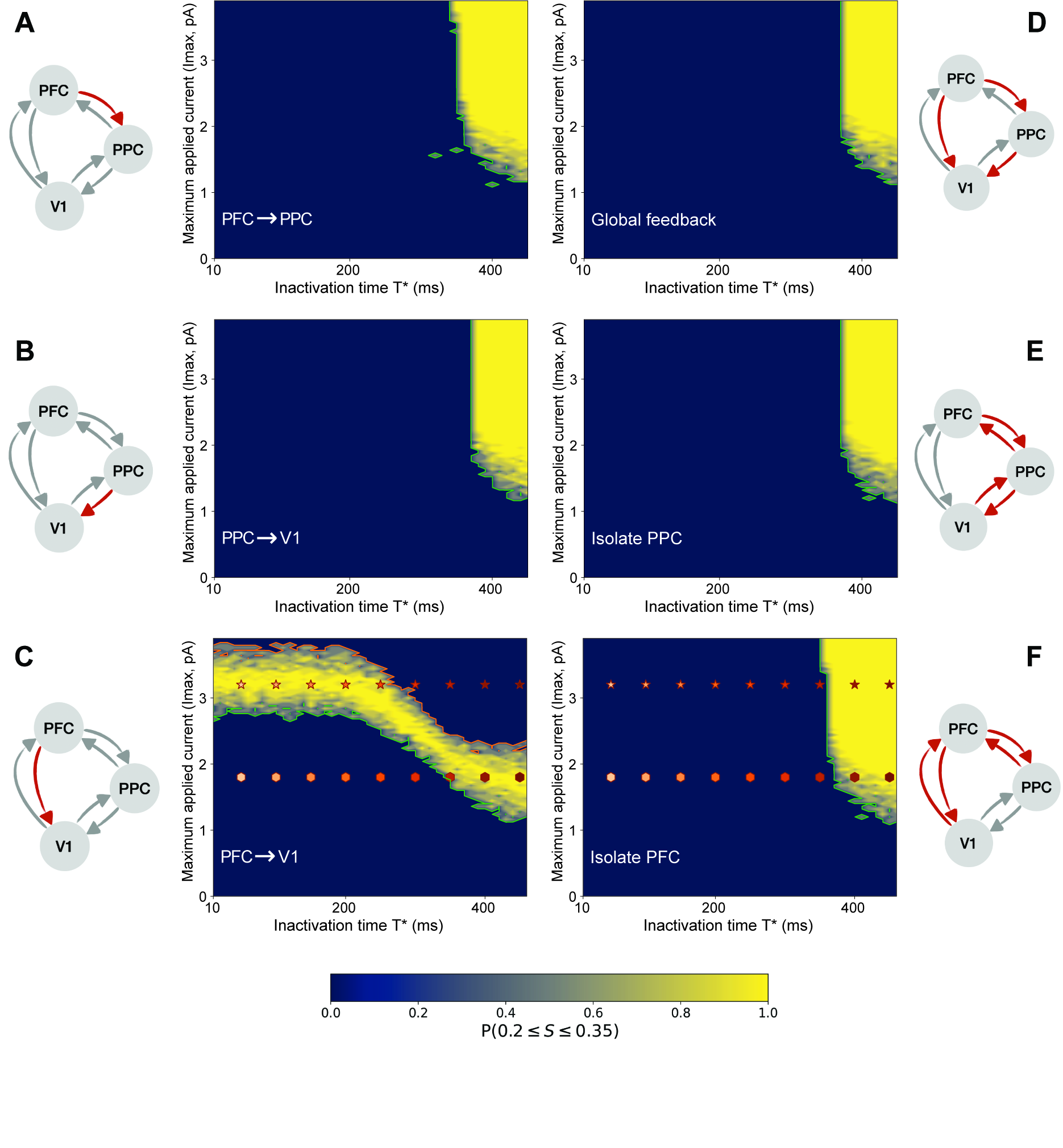
